# ATP-Independent Water-Soluble Luciferins Enable Non-Invasive High-Speed Video-Rate Bioluminescence Imaging of Mice

**DOI:** 10.1101/2024.04.30.591933

**Authors:** Xiaodong Tian, Yiyu Zhang, Hui-wang Ai

## Abstract

NanoLuc luciferase and its derivatives are attractive bioluminescent reporters recognized for their efficient photon production and ATP independence. However, utilizing them for *in vivo* imaging poses notable challenges. Low substrate solubility has been a prominent problem, limiting *in vivo* brightness, while substrate instability hampers consistent results and handling. To address these issues, we developed a range of caged PEGylated luciferins with improved stability and water solubility of up to 25 mM, resulting in substantial bioluminescence increases in mouse models. This advancement has created the brightest and most sensitive luciferase-luciferin combination, enabling high-speed video-rate imaging of freely moving mice with brain-expressed luciferase. Furthermore, we developed a bioluminescent Ca^2+^ indicator with exceptional sensitivity to physiological Ca^2+^ changes and paired it with a new substrate to showcase non-invasive, video-rate imaging of Ca^2+^ activity in a defined brain region in awake mice. These innovative substrates and the Ca^2+^ indicator are poised to become invaluable resources for biological and biomedical fields.

---

Bioluminescence imaging (BLI) is a powerful, non-invasive method for monitoring biological processes in animal models.^1,2^ Unlike fluorescence imaging, it avoids issues like photobleaching and phototoxicity.^3^ BLI, utilizing luciferase-catalyzed oxidation of luciferins for photon production, enables sensitive signal monitoring in deep tissues with high signal-to-background ratios. Compared to alternative *in vivo* imaging modalities such as magnetic resonance imaging (MRI) and positron emission tomography (PET), BLI offers superior spatiotemporal resolution, cost-effectiveness, convenience, and the absence of radioactive contrast agents.^4,5^ When combined with bioluminescent indicators, BLI can track specific bioactivities, making it a popular imaging technique for both basic and preclinical research.^6,7^

Firefly luciferase (FLuc), which utilizes D-luciferin as its substrate, is a widely adopted bioluminescent reporter due to its long-wavelength light emission and the substrate’s chemical stability and water solubility. Research has developed FLuc and D-luciferin derivatives offering enhanced properties such as higher brightness, more red-shifted emission, and orthogonal reactivity.^1,8-11^ Of particular note is the Akaluc luciferase coupled with the AkaLumine luciferin (also known as TokeOni), recognized as a leading benchmark for deep-tissue sensitivity.^12^ However, these luciferases and their derived bioluminescent indicators inherently depend on ATP, a crucial energy source and signaling molecule, for their operation.^7,13^

Conversely, the oxidation of coelenterazine (CTZ) luciferin by marine luciferases is ATP-independent. Promega introduced NanoLuc, a marine luciferase mutant, known for its high photon production with the synthetic luciferin named furimazine (Fz), small molecular size, remarkable enzyme stability, and flexibility for split and domain insertion.^14,15^ Despite its popularity, NanoLuc faces challenges for *in vivo* BLI, such as limited tissue penetration of the emitted blue photons, low substrate stability and solubility, and inadequate luciferin entry to the brain.^13,16,17^ Recent studies partially addressed these concerns by developing additional CTZ analogs and NanoLuc mutants,^18-20^ or by fusing luciferases to long-wavelength-emitting fluorescent proteins (FPs) for redder emission.^18,21-23^ Furthermore, NanoLuc and its derived luciferases have been effectively transformed into bioluminescent indicators, enabling successful imaging of Ca^2+^ and K^+^ dynamics in live mice.^23-25^ Despite advancements, there is a pressing need for further research to systematically improve the *in vivo* performance of NanoLuc-derived luciferase-luciferin pairs and fully realize the potential of BLI in living animals.

Here, we present a PEGylation method to enhance the stability and water solubility of marine luciferase substrates. We showcased the versatility of this approach using three luciferins, resulting in a series of new prosubstrates with remarkably increased solubility. Given the challenges of brain BLI due to the blood-brain barrier (BBB) limiting luciferin entry and the need for capturing rapid neuronal activity,^17^ we focused on characterizing these prosubstrates using mouse models expressing luciferases in the brain. We observed significant bioluminescence enhancements with these water-soluble luciferins that are deliverable at a safe concentration of 25 mM. Notably, we identified the brightest luciferase-luciferin combination, which surpassed the brightness of Akaluc and AkaLumine, allowing high-speed video-rate imaging of freely moving mice with brain-expressed luciferase. Furthermore, in a mouse model with liver luciferase expression, we validated the substantial enhancement in brightness achieved using one of the novel luciferins. Furthermore, we developed a significantly enhanced bioluminescent Ca^2+^ indicator that effectively responds to physiological Ca^2+^ dynamics. We demonstrate non-invasive video-rate Ca^2+^ imaging of footshock-induced neuronal activation in awake mice by using a water-soluble luciferin and expressing the Ca^2+^ indicator in the basolateral amygdala (BLA) region of the brain.

## RESULTS

### PEGylation of DTZ and solubility and stability assessment

We previously developed a NanoLuc variant, teLuc, that produced bright bioluminescence peaking around 500 nm when combined with a DTZ substrate (**Fig. 1a**).^18^ More recently, we refined DTZ to create a prosubstrate, ETZ, which could be administered in higher doses and activated *in vivo* by nonspecific esterase, enhancing *in vivo* bioluminescence.^23^ In parallel, other researchers introduced luciferin variants such as HFz and FFz, improving water solubility and brightness in non-brain tissues.^26^ However, the solubility improvements from these prior studies were modest, and high *in vivo* brightness still required the use of high doses of organic cosolvents or surfactants, which could potentially induce organ toxicity.^17,27^

**Fig. 1.**
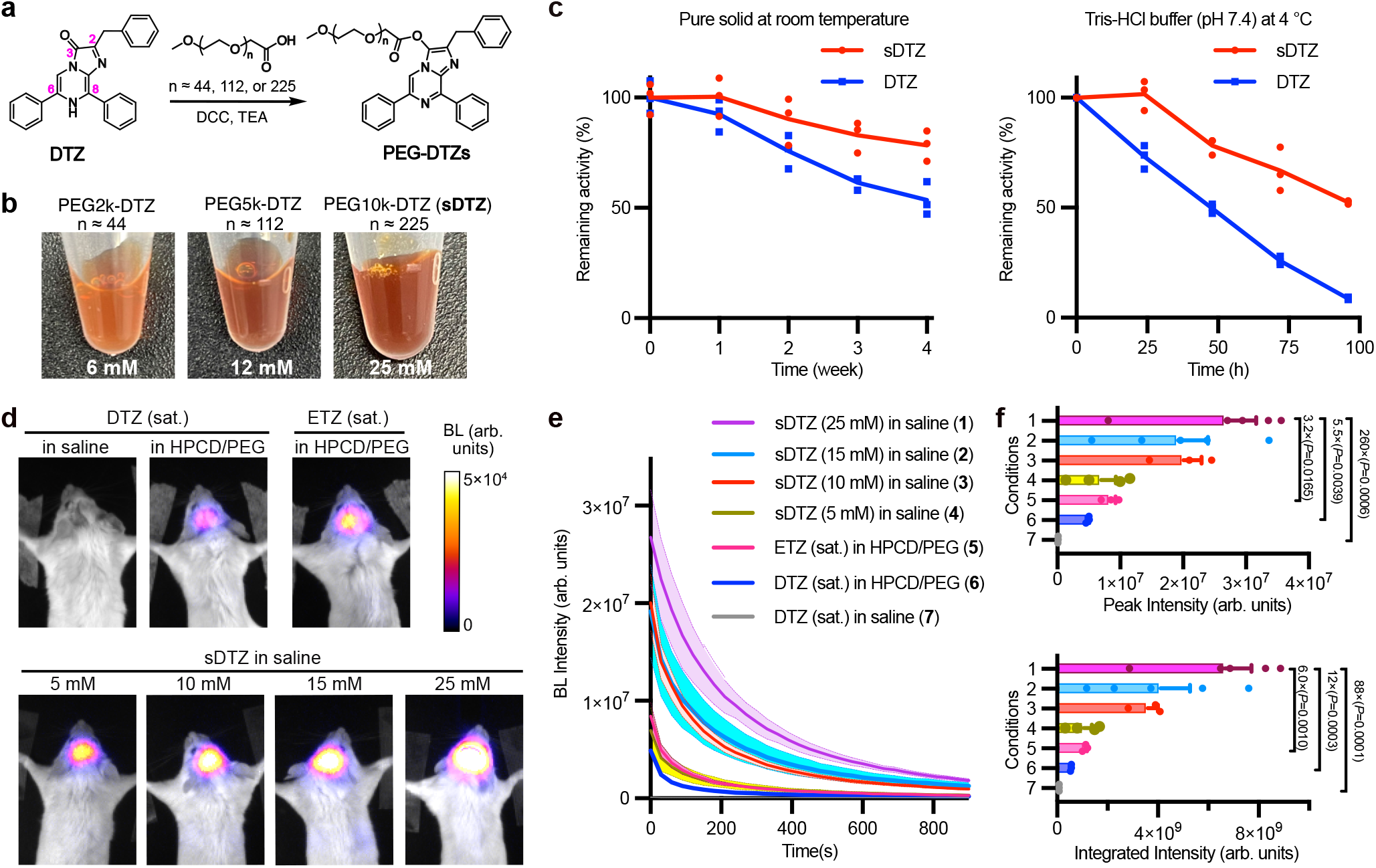
Synthesis of water-soluble DTZ derivatives and evaluation of solubility, stability, and sensitivity for *in vivo* bioluminescence brain imaging. (**a**) Synthetic route to modify DTZ with three lengths of polyethylene glycol (PEG) chains (PEG 2k, 5k, or 10k) via the C3 carbonyl group of DTZ. (**b**) Photos of PEGylated DTZ variants dissolved in normal saline at the indicated near-saturation concentrations. (**c**) Comparison of DTZ and sDTZ for solid-state stability at room temperature or stability in aqueous buffer (pH 7.4) at 4 °C. n = 3 technical replicates. (**d**) Representative bioluminescence images of live mice with hippocampal BREP luciferase expression upon the tail vein delivery of the luciferins at the indicated dose and buffer conditions. Images with peak bioluminescence intensities were presented in pseudocolor overlaid on corresponding brightfield images. (**e**) Quantification of bioluminescence intensity over time for each group in panel d. (**f**) Comparison of peak bioluminescence intensity (top) and bioluminescence integrated over 15 min (bottom). The numbered conditions refer to the groups presented in panel e. *P* values were determined by ordinary one-way ANOVA followed by Dunnett’s multiple comparisons test. Data are presented as mean ± s.e.m. (in panels d-f, n = 4 mice for 5 mM sDTZ, n = 5 mice for 15 and 25 mM sDTZ, and n = 3 mice for other groups). BL, bioluminescence. Arb. units, arbitrary units.

We aimed to significantly enhance the water solubility of marine luciferase substrates, enabling high-dose delivery via aqueous solutions. Polyethylene glycol (PEG) conjugation is a commonly employed method for improving the solubility of hydrophobic drugs.^28^ Using DTZ as our model compound, we investigated PEGylation at the C3 carbonyl group of the DTZ imidazopyrazine ring (**Fig. 1a**). We expected PEGylation at this site to create protected substrates resistant to oxidation, but upon injection into animals, nonspecific esterase would remove the PEGylation, releasing free DTZ luciferin.^23^

By conducting condensation reactions of DTZ with methoxy PEG carboxylic acids of varying molecular weights (∽2000, 5000, and 10000 Da), we obtained the compounds (PEG2k-DTZ, PEG5k-DTZ, and PEG10k-DTZ) and determined their maximum solubility in normal saline to be 6, 12, and 25 mM, respectively (**Fig. 1b**). Given that PEG10k-DTZ exhibited the highest solubility, we renamed it sDTZ and focused our subsequent experiments on this compound.

Next, we evaluated the chemical stability of sDTZ and found it to be more stable than DTZ under all conditions. As a solid, sDTZ retained stability for about a week at room temperature, gradually degrading thereafter but maintaining roughly 80% activity after four weeks (**Fig. 1c**). The slow degradation of sDTZ was likely due to the gradual removal of PEGylation, prompted by atmospheric moisture absorption. Moreover, when dissolved in Tris-HCl (pH 7.4) buffer, sDTZ remained stable for a day at 4 °C and for a week at either −20 or −80 °C (**Fig. 1c**).

### Brightness enhancement for brain and liver imaging, and toxicity evaluation

We subsequently evaluated sDTZ for enhancing BLI brightness using mouse models. In a prior study, we developed an optimized fusion protein named BREP by combining teLuc with the red fluorescent protein (RFP) mScarlet-I.^23^ BREP emits about 60% of its total emission above 600 nm, positioning it as a highly effective luciferase for deep-tissue BLI. We generated adeno-associated viruses (AAVs) harboring the BREP gene under the control of the human synapsin I (hSyn) promoter and introduced them into the hippocampal neurons of mice through stereotactic injection. Two to three weeks later, the mice with BREP expressed in the brain were examined under anesthesia in a dark box with an EMCCD camera after receiving 100 μL of luciferin solutions via the tail vein.

Experimenting with four different sDTZ concentrations revealed that higher luciferin concentrations led to increased bioluminescence intensity (**Fig. 1d**). The brightest signal occurred with a near-saturation sDTZ concentration (25 mM) in normal saline. The signal decreased over time and around 7% of the peak intensity was retained after 15 min (**Fig. 1e**). In contrast, DTZ saturated in saline had a peak intensity 260-fold lower and an 15-min integrated intensity 88-fold lower (**Fig. 1f**). Strong bioluminescence was observed when DTZ and ETZ were dissolved in a formulation containing 25% (w/v) 2-hydroxypropyl-β-cyclodextrin (HPCD) and 20% (v/v) PEG-400,^23^ which allowed the maximal final concentrations of DTZ and ETZ to be approximately 2.5 mM and 6.8 mM, respectively. However, the sDTZ in saline group still had a peak intensity 5.5-fold and 3.2-fold higher than DTZ and ETZ, respectively. In addition, the integrated bioluminescence signal over 15 min for the sDTZ in saline group was approximately 12-fold and 6-fold higher than the DTZ and ETZ in HPCD/PEG groups, respectively (**Fig. 1f**). Collectively, the results indicate that sDTZ enables efficient *in vivo* luciferin delivery using normal saline, significantly enhancing sensitivity for brain BLI in mice.

To explore bioluminescence improvement in different tissues, we introduced BREP luciferase into mouse livers using AAVs with a hepatocyte-specific thyroxine-binding globulin (TBG) promoter.^20,29^ Mice were administered different concentrations of sDTZ before imaging. The bioluminescence intensity increased with higher sDTZ concentrations, exceeding that of DTZ in saline by at least two orders of magnitude. The 25 mM sDTZ concentration resulted in the highest intensity, with nearly a 1000-fold peak intensity increase and a 482-fold integrated intensity increase over the first 15 min. These findings align with brain imaging results, confirming substantial sensitivity enhancement by sDTZ in various tissue types.

After determining 25 mM sDTZ as optimal for achieving maximum sensitivity, we assessed organ toxicity with five consecutive daily injections. At the end of the experiment, mice were sacrificed, and organs were extracted and examined with hematoxylin and eosin (H&E) staining. No apparent organ toxicity was found, suggesting sDTZ’s safety for animal imaging.

### PEGylation of alternative luciferins and brain brightness evaluation

After achieving success with sDTZ, we extended the PEGylation method to other luciferins. Specifically, we focused on two crucial ones: furimazine (Fz),^14^ the original NanoLuc substrate, and CFz,^17^ a recently developed NanoLuc substrate with improved blood-brain barrier (BBB) penetration but reduced water solubility. We introduced a methoxy PEG chain (MW ∽10,000) to create sFz and sCFz, respectively (**Fig. 2a**). To assess sFz and sCFz for in vivo BLI, we used AAVs to express the Antares luciferase,^21^ a fusion of NanoLuc to two copies of CyOFP1, in the hippocampal region of the mouse brain. We injected sFz or sCFz (25 mM in normal saline) via the tail vein and compared results with Fz or CFz saturated in the HPCD/PEG buffer mentioned earlier, as well as normal saline. Both sFz and sCFz significantly enhanced brightness (**Fig. 2b-f**). Particularly, sCFz led to a 179-fold and 27-fold peak bioluminescence increase compared to CFz in normal saline and the HPCD/PEG buffer, respectively. Moreover, the sCFz group showed approximately 5-6-fold brighter signals than sFz (**Fig. 2ef**), consistent with CFz’s better BBB permeability than Fz.^17^ These findings indicate that PEGylation can improve solubility and greatly enhance *in vivo* BLI sensitivity across different luciferins.

**Fig. 2.**
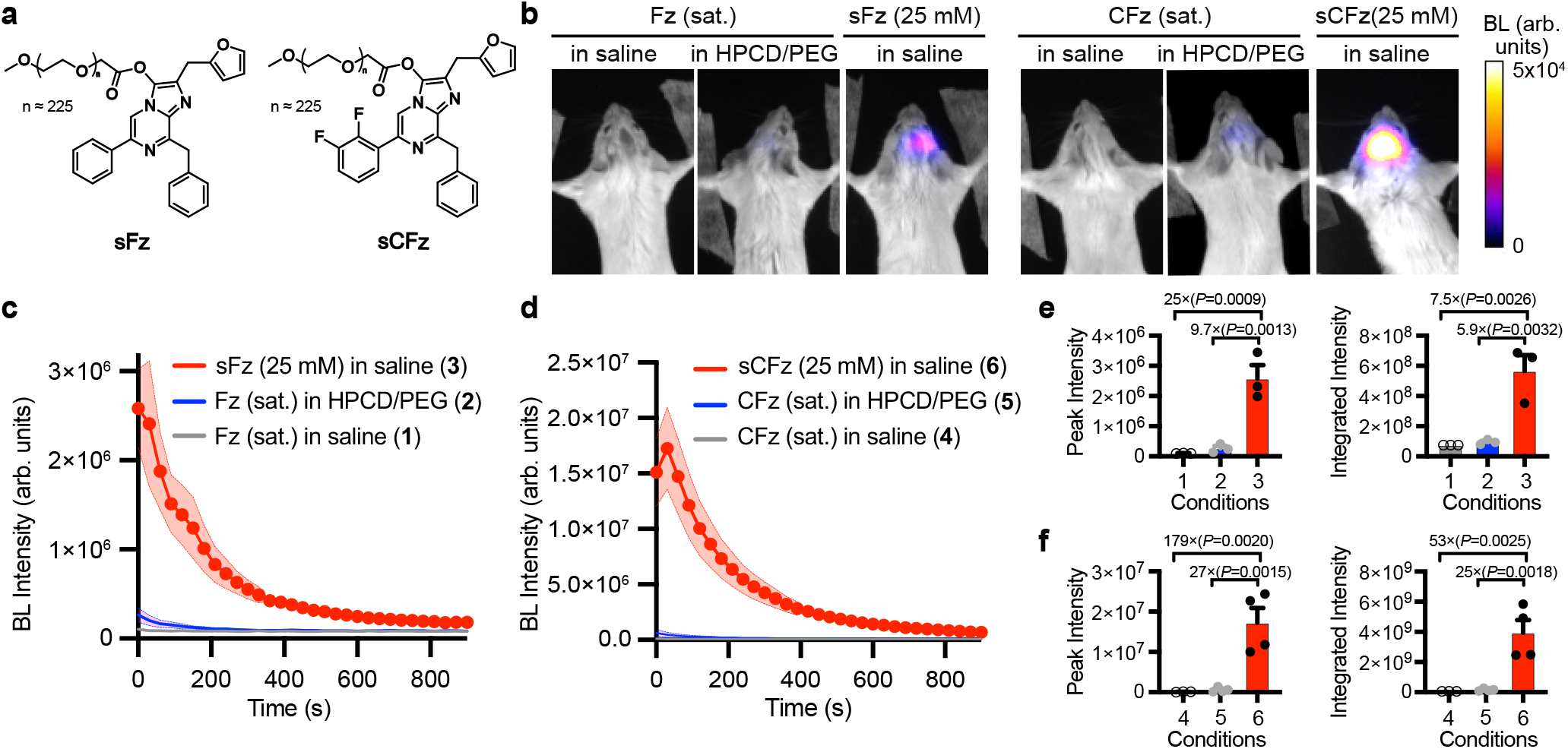
Evaluation of water-soluble Fz and CFz variants, sFz and sCFz, paired with Antares for *in vivo* bioluminescence brain imaging. (**a**) Chemical structures of sFz and sCFz. (**b**) Representative bioluminescence images of live mice with hippocampal Antares luciferase expression upon the tail vein delivery of the luciferins at the indicated dose and buffer conditions. Images with peak bioluminescence intensities were presented in pseudocolor overlaid on corresponding brightfield images. (**c**) Quantification of bioluminescence intensity over time for mice in the Fz and sFz groups. (**d**) Quantification of bioluminescence intensity over time for mice in the CFz and sCFz groups. (**e**) Comparison of peak bioluminescence intensity (left) and bioluminescence integrated over time (right) for the Fz and sFz groups. (**f**) Comparison of peak bioluminescence intensity (left) and bioluminescence integrated over time (right) for the CFz and sCFz groups. In panels e and f, the numbered conditions refer to the groups presented in panels c and d; *P* values were determined by ordinary one-way ANOVA followed by Dunnett’s multiple comparisons test. Data are presented as mean ± s.e.m. (in panels b-f, n = 4 mice for CFz in HPCD/PEG and sCFz groups, and n = 3 mice for other groups). BL, bioluminescence. Arb. units, arbitrary units.

### Identification of the optimal luciferase-luciferin combination and high-speed video-rate brain imaging of freely moving mice

Based on our data, both BREP-sDTZ and Antares-sCFz combinations displayed bright bioluminescence, with BREP-sDTZ being roughly twice as bright as Antares-sCFz (**Figs. 1f** and **2f**). Considering the cross-reactivity of teLuc in BREP towards NanoLuc substrates, we further examined the bioluminescence of mice with hippocampal BREP expression after delivering sCFz (**Fig. 3a**). The BREP-sCFz mice exhibited strong bioluminescence, with intensity below the BREP-sDTZ mice (**Fig. 3bc**). Although the sample size was inadequate to statistically distinguish the peak intensities, BREP-sCFz displayed a quicker bioluminescence decay and lower 15-min integrated bioluminescence that is statistically significant compared to BREP-sDTZ (**Fig. 3bc**). These findings indicate that the BREP-sDTZ combination exhibits the most promise among all ATP-independent luciferase-luciferin pairs.

**Fig. 3.**
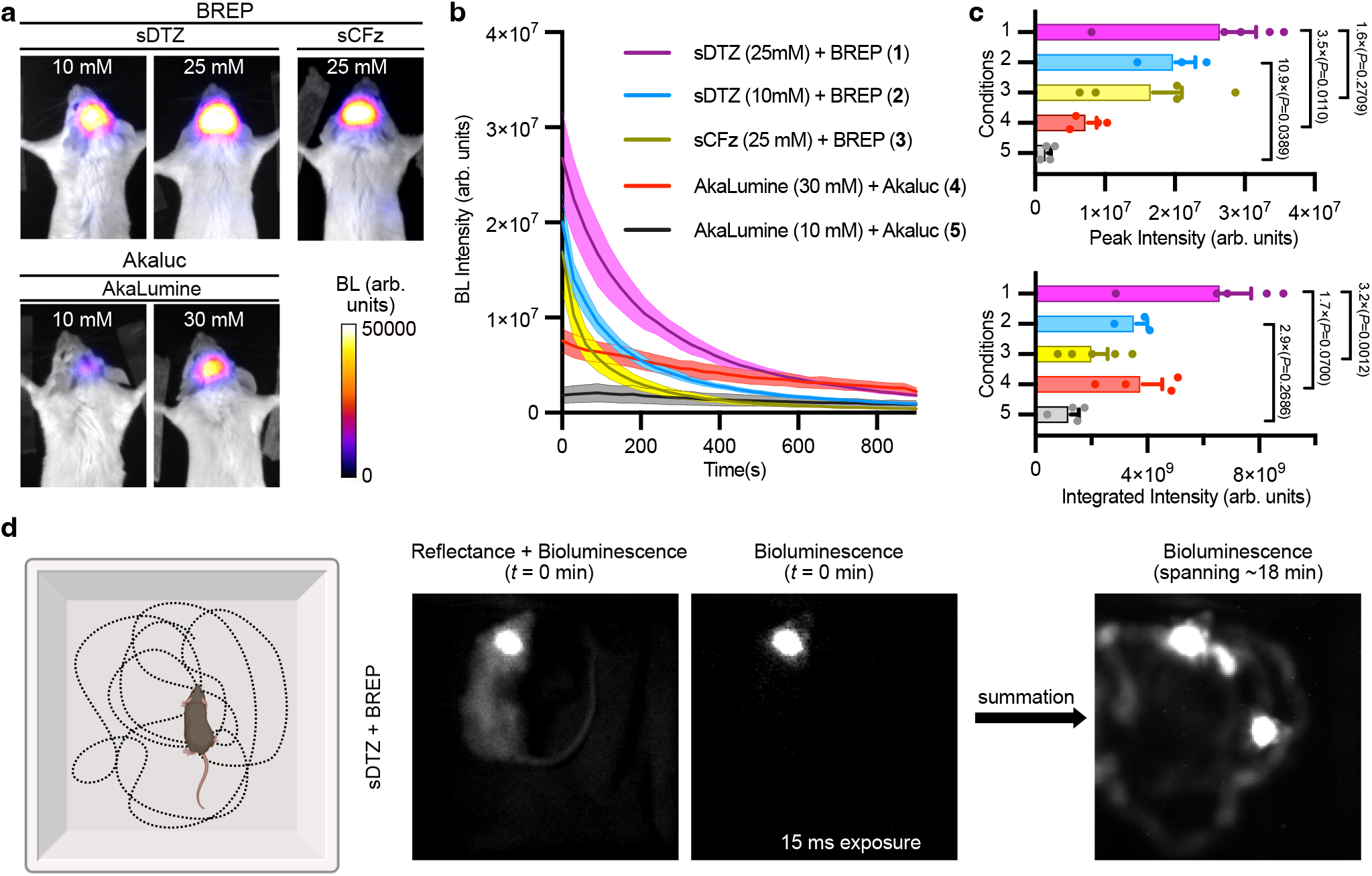
Further brightness comparison and high-frame video-rate *in vivo* bioluminescence brain imaging using BREP and sDTZ. (**a**) Representative bioluminescence images of live mice with hippocampal luciferase expression upon the tail vein delivery of the indicated luciferins at the indicated dose using normal saline. Comparisons were made between BREP (paired with sDTZ or sCFz) and Akaluc (paired with Akalumine). Images with peak bioluminescence intensities were presented in pseudocolor overlaid on corresponding brightfield images. (**b**) Quantification of bioluminescence intensity over time among different groups. (**c**) Comparison of peak bioluminescence intensity (top) and bioluminescence integrated over time (bottom). The numbered conditions refer to the groups presented in panel b. *P* values were determined by ordinary one-way ANOVA followed by Tukey’s multiple comparison tests. Data are presented as mean values ± s.e.m. (n = 3 mice for BREP and 10 mM sDTZ, n = 4 mice for Akaluc and 10 or 30 mM AkaLumine, and n = 5 mice for all other groups). The same data for BREP and sDTZ in panels a-c are also presented in Fig. 1d-f. (**d**) High frame-rate imaging of a mouse with hippocampal BREP luciferase expression. The mouse was injected with sDTZ via the tail vein right before imaging. Brightfield and bioluminescent images (15-ms exposure time for each) were alternately acquired using an EMCCD camera. The first frame (at *t* = 0 min) was shown. A series of 280 frames of bioluminescent images at each time point (*t* = 0, 3, 6, 12 and 18 min) were acquired, and the total 1,400 frames were combined to create the summation image. BL, bioluminescence. Arb. units, arbitrary units.

In addition, we compared BREP-sDTZ with the ATP-dependent luciferase-luciferin pair, Akaluc-AkaLumine.^12^ All luciferins were administered intravenously in normal saline to mice with hippocampal luciferase expression. At 10 mM luciferin concentrations, the peak intensities of BREP mice were approximately 11 times brighter than Akaluc mice (**Fig. 3a-c**). With 25 mM sDTZ and 30 mM AkaLumine, BREP remained 3.5 times brighter than Akaluc. Despite displaying a faster bioluminescence decay, BREP-sDTZ consistently exhibited greater brightness in the initial 10 min, indicating its suitability for imaging applications demanding high sensitivity (**Fig. 3b**).

Given the excellent performance of sDTZ, we conducted high-speed video-rate BLI of freely moving mice expressing BREP in the hippocampus (**Fig. 3d**). The bioluminescence glow from the brain was so bright and even visible in brightfield imaging at the beginning of the experiment. By alternating between brightfield and bioluminescence imaging, we successfully captured mouse movement and brain bioluminescence for over 18 min using a 15-ms exposure. Previous research has demonstrated the capability to monitor Akaluc and Antares signals in moving mouse brains using AkaLumine or CFz.^12,17^ Despite variations in protocols and equipment, our study achieved enhanced temporal resolution by utilizing BREP-sDTZ, even in unshaved mice.

### Development of a high-quality bioluminescent Ca^2+^ indicator and non-invasive video-rate Ca^2+^ imaging in awake mice

Having confirmed the enhanced *in vivo* BLI sensitivity provided by sDTZ, we proceeded to investigate its potential to facilitate functional imaging. Calcium (Ca^2+^) serves as an indicator of neuronal activity, and bioluminescent imaging of Ca^2+^ offers a non-invasive method for monitoring the activity of specific neuronal populations in animals.^17,23^ Several bioluminescent Ca^2+^ indicators have been developed to date.^22,24,30-33^ Our previously reported indicator, BRIC (bioluminescent red indicator for Ca^2+^), stands out due to its superior brightness, response magnitude, and red-shifted emission, making it an excellent choice for *in vivo* BLI.^23^

Here we first enhanced the responsiveness of BRIC to physiological Ca^2+^ fluctuations. To achieve this, we conducted random mutagenesis on BRIC and screened the resulting libraries using 65 nM and 1.35 μM of free Ca^2+^, which differs from previous screening approaches that used 0 and 39 μM of free Ca^2+^.^23,24^ Through three rounds of directed evolution, we successfully generated an enhanced BRIC variant (eBRIC) with five mutations (**Fig. 4a**). The response of the variant between 65 nM and 1.35 μM of Ca^2+^ increased from 2-fold to 4.5-fold, while the overall Ca^2+^-induced change between 0 and 39 μM of free Ca^2+^ increased from 6.5-fold to 17-fold (**Fig. 4b**). The apparent dissociation constant (*K*_d_) also increased from 133 nM to 2.3 μM (**Fig. 4c**). We further conducted a comparative analysis of eBRIC and BRIC for imaging Ca^2+^ dynamics in HeLa cells and primary neurons. In eBRIC-expressing HeLa cells, the stimulation with 20 μM histamine resulted in robust Ca^2+^ oscillations that were nearly 4-fold higher than BRIC under the same conditions (**Fig. 4d**). In primary neurons, high K^+^ depolarization induced a 7.5-fold increase in eBRIC bioluminescence, resulting in a 5.8-fold increase in responsiveness (ΔBL/BL_0_) from BRIC (**Fig. 4e**). Notably, two of the five mutations in eBRIC are located within the Ca^2+^-binding calmodulin (CaM) and M13 elements (**Fig. 4a**). Based on our ColabFold model,^34^ the N515D mutation is at the CaM-M13 interaction interface, while the D474E mutation affects an H-bond near the interface. These mutations are likely crucial for the enhanced properties of eBRIC.

**Fig. 4:**
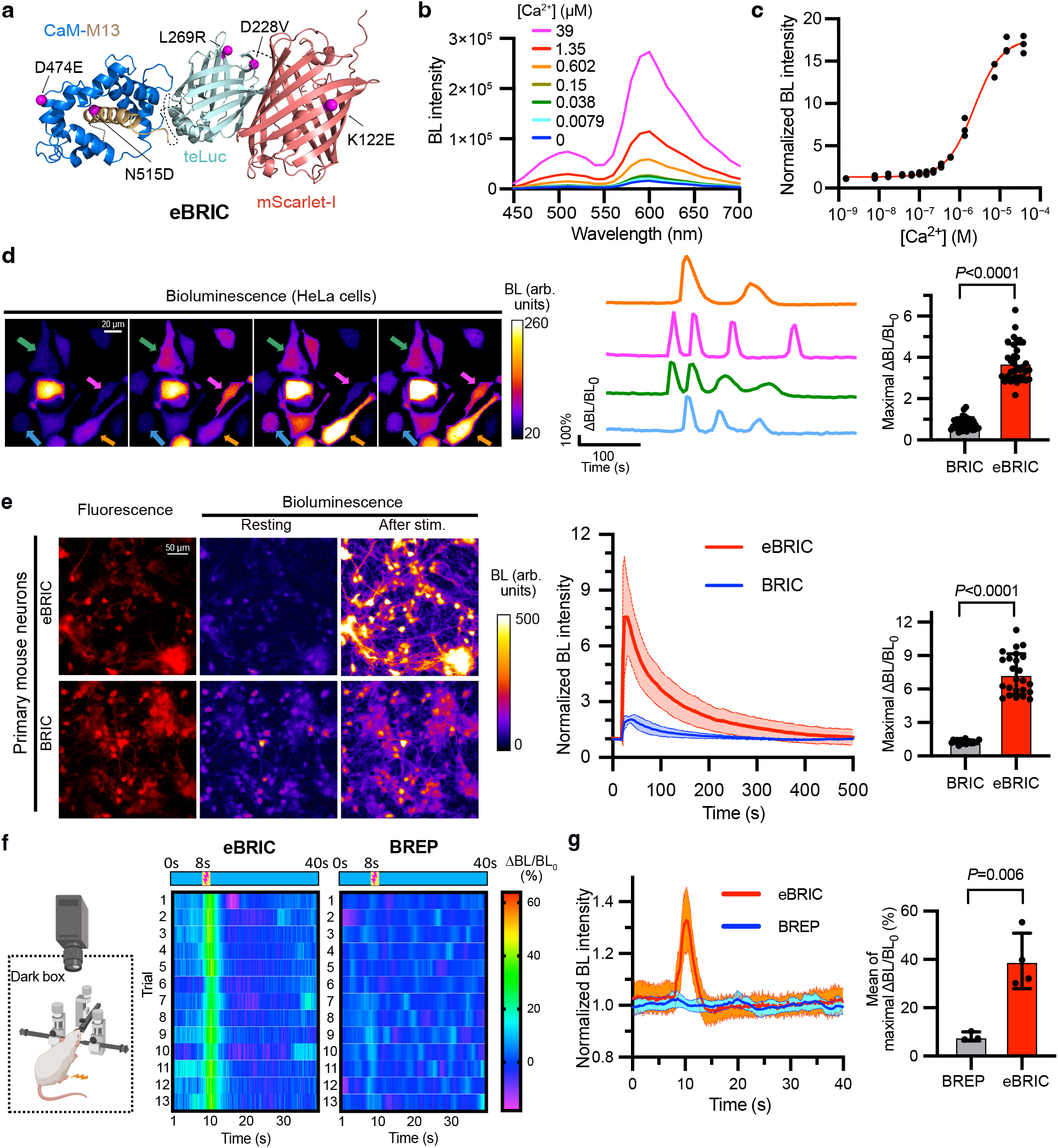
Engineering of eBRIC, *in vitro* characterization, and the application of eBRIC and sDTZ for video-rate BLI of Ca^2+^ dynamics in the awake mouse brain. (**a**) Domain arrangement of eBRIC with five mutations from BRIC highlighted as magenta balls. The model structure was created using ColabFold v1.5.5. The CaM (calmodulin), M13, teLuc, and mScarlet-I domains are colored in marine blue, golden yellow, pale cyan, and salmon pink, respectively. (**b**) Bioluminescence spectra of eBRIC in the presence of DTZ and varying concentrations of free Ca^2+^. (**c**) Ca^2+^-dependency of eBRIC bioluminescence at 600 nm (n = 3 technical replicates). A four-parameter Hill equation was used to fit the data to determine the dissociation constant (*K*_d_ = 2.3 μM and Hill coefficient = 1.3). (**d**) **Left:** Representative pseudocolored bioluminescence images (left) of histamine-induced Ca^2+^ dynamics in HeLa cells. Scale bar, 20 μm. **Middle:** Intensity traces for individual cells. The color of the traces corresponds to the colors of the arrows used to highlight individual cells in the left images. The baselines of the traces were corrected for monoexponential decay resulting from substrate consumption. **Right:** Parallel comparison between eBRIC and BRIC in terms of the maximal changes (ΔBL/BL_0_) of individual cells induced with histamine. Data are expressed as mean ± s.d. (n= 33 cells for eBRIC and 34 cells for BRIC), with the *P* value derived from unpaired two-tailed *t*-tests. (**e**) **Left:** Representative fluorescence and bioluminescence images of primary mouse neurons expressing eBRIC or BRIC in response to high K^+^ (30 mM) depolarization. Scale bar, 50 μm. **Middle:** Quantification of the responses of eBRIC- or BRIC-expressing neurons. **Right:** Comparison of the maximal changes (ΔBL/BL_0_) of individual cells. Data are expressed as mean ± s.d. (n = 24 cells for eBRIC and 17 for BRIC), with the *P* value determined through unpaired two-tailed *t*-tests. (**f**) **Left:** Scheme of BLI of head-fixed awake mice. eBRIC expression in the basolateral amygdala (BLA) region was achieved using AAV vectors. Prior to imaging, sDTZ was administered to mice via the tail vein. Footshocks were then employed as a means of stimulation. Images were acquired with a 100-ms exposure time without any intervals. **Right:** Heatmap illustrating bioluminescence intensity responses of a representative mouse expressing either eBRIC or BREP (negative control) following 13 consecutive footshock stimulation trials. (**g**) **Left:** Quantification of intensity changes of eBRIC- and BREP-expressing mice in response to footshock. Data is presented as mean ± s.d. and based on images of 4 mice for eBRIC and 3 mice for BREP, each consisting of 13 trials. **Right:** Comparison of the eBRIC and BREP groups by averaging the responses of individual mice over 13 trials. Data are expressed as mean ± s.d. (n = 4 mice for eBRIC and 3 mice for BREP), with the *P* value derived from unpaired two-tailed *t*-tests.

We paired sDTZ with eBRIC for non-invasive imaging of neuronal activity *in vivo* (**Fig. 4f**). The basolateral amygdala (BLA) region plays a crucial role in generating fear-associated behaviors.^35^ We produced adeno-associated viruses (AAVs) carrying the eBRIC gene and introduced them into the BLA regions in mice. Post eBRIC expression, we administered sDTZ and applied footshock stimuli to activate the BLA neurons. We noticed a significant and consistent increase in bioluminescence in the eBRIC group (**Fig. 4g**). As a control, we included Ca^2+^-insensitive BREP, which exhibited minimal to no changes in bioluminescence under electric footshock stimulation. Compared to prior similar experiments with BRIC,^23^ the current study achieved a superior temporal resolution, enhancing it from 1 second to 100 milliseconds, along with an approximately twofold increase in response magnitude. These findings confirm that sDTZ and eBRIC serve as powerful tools for tracking the activity of neuronal populations with improved temporal resolution and sensitivity.

## DISCUSSION

The primary contribution of this work is the development of a PEGylation method for modifying marine luciferase substrates. By attaching PEG chains to these substrates, their solubility and stability were greatly enhanced, allowing for effective *in vivo* luciferin administration using a simple and non-toxic normal saline solution. The resulting luciferase-luciferin pairs exhibited notable increases in brightness, surpassing established benchmarks such as Akaluc and Antares.^12,21^ This research tackles critical obstacles in the field, achieving optimal sensitivity with ATP-independent luciferase reporters and enabling high-speed video-rate imaging of freely moving animals. In addition, eliminating organic cosolvents or surfactants from the luciferin delivery solution addresses the *in vivo* toxicity problem,^17^ leading to enhanced biocompatibility of BLI. Furthermore, the extension of the PEGylation technology to diverse luciferins represents a notable progression. Demonstrating the adaptability of PEGylation with various luciferin substrates in this research widens the range of applications and research possibilities, highlighting the potential for personalized solutions tailored to specific imaging needs.

Our validation experiments were centered on brain imaging, considering the challenges posed by the BBB and the essential requirement for high temporal resolution to capture rapid neuronal activities. The identification of the most luminous luciferase-luciferin combination opens up avenues for accurate and sensitive imaging of brain targets in live organisms, showcasing the potential for exploring various questions spanning from neurobiology to cancer research.

Furthermore, we developed an enhanced bioluminescent Ca^2+^ indicator (eBRIC) by using a physiological Ca^2+^ concentration range during library screening. As BRIC has been set as a benchmark for bioluminescent Ca^2+^ indicators,^23^ eBRIC signifies a substantial advancement beyond BRIC. Indeed, the combination of eBRIC with the novel luciferin substrate expands the capability of functional neuronal imaging. The successful demonstration of non-invasive, video-rate imaging of Ca^2+^ activity in a defined brain region in awake mice represents a significant milestone. The improved temporal resolution and response magnitude offered by eBRIC and sDTZ underscore their potential as powerful tools for investigating dynamic processes in living organisms.

This study further illustrates that the application of PEGylated luciferins extends to non-brain tissues. The use of sDTZ allowed for high-sensitivity BLI of the liver, indicating that the novel tools can be applied to the study of various organ-specific processes, disease models, and therapeutic interventions in diverse tissue types.

In summary, this work has yielded novel caged PEGylated luciferins and a significantly improved high-performance bioluminescent Ca^2+^ indicator, serving as invaluable assets for furthering our comprehension of biological processes and disease mechanisms. These new tools provide researchers with unparalleled sensitivity, resolution, and versatility in investigating complex biological systems.

## METHODS

### Chemical synthesis and characterization

Methoxy PEG acetic acids with molecular weights of ∽2,000, ∽5,000, and ∽10,000 Da were procured from JenKem Technology. DTZ, Fz, and CFz were synthesized following established procedures^14,17,18^ and purified using a Waters Prep 150/SQ Detector 2 LC-MS Purification System with MassLynx software (Version 4.2) and an XBridge BEH Amide/Phenyl OBD Prep Column (130 Å, 5 μm, 30 mm × 150 mm). For the conjugation reaction, methoxy PEG acetic acid (0.02 mmol, 1 equiv.) and DCC (12.4 mg, 0.06 mmol, 3 equiv.) were combined in a dried and argon-purged two-neck 50 mL round-bottom flask. A mixture of anhydrous dichloromethane (10 mL) and triethylamine (4.4 μL, 0.06 mmol, 3 equiv.) was slowly added while stirring at room temperature for 20 minutes. Then, DTZ, Fz, or CFz (0.024 mmol, 1.2 equiv.) was swiftly introduced into the reaction mixture. The reaction was stirred under argon for an additional hour and monitored by TLC (DCM/methanol = 10:1). Upon completion, the reaction mixture was concentrated *in vacuo*, and the residue was washed with ethyl ether and purified by silica gel chromatography (DCM/methanol = 15:1, v/v). The resulting compound was collected, concentrated, dissolved in 5 mL ddH_2_O, filtered through VWR Grade 415 qualitative filter paper, and lyophilized to obtain sDTZ, sFz, or sCFz as solids using a 12-port Labconco freeze dryer with an Edwards RV3 vacuum pump. ^1^H-NMR spectra were recorded on a Bruker Avance III 600 MHz NMR spectrometer with Bruker TopSpin IconNMR software (Version 3.5pl4) and analyzed using MestReNova software (Version 12.0.3). Residual solvent peaks were referenced at δ=4.79 ppm (D_2_O) or 3.31 ppm (CD_3_OD).

### Chemical stability evaluation

The stabilities of DTZ and sDTZ were evaluated in two forms: as 100 μM solutions in Tris-HCl buffer (pH 7.4) at temperatures of 4, −20, and −80 °C, and as powders stored at room temperature in aluminum foil-wrapped microcentrifuge tubes to shield from light exposure. The integrity of the chemicals was assessed using a bioluminescence assay with purified BREP luciferase. For the assays, solid powders were dissolved in Tris-HCl buffer (pH 7.4) to obtain 100 μM solutions. DTZ solutions were directly used, while sDTZ solutions underwent pretreatment with porcine liver esterase (MilliporeSigma, Cat. # E3019) by incubating 100 μL of the sDTZ solution with 1 μL of 50 units/mL enzyme for 15 minutes at room temperature. Subsequently, DTZ or esterase-treated sDTZ were mixed with purified BREP protein to achieve final concentrations of 50 μM luciferin and 10 nM enzyme. Bioluminescence spectra were recorded using a BMG Labtech CLARIOstar Plus microplate reader controlled by BMG Labtech Reader Software (Version 5.70 R2), with results automatically exported to BMG Labtech MARS Data Analysis Software (Version 3.42 R5).

### Toxicity assessment in mice

sDTZ (25 mM) was dissolved in normal saline, and a daily intraperitoneal injection of 100 μL of the solution was administered to C57BL/6J mice for five consecutive days. Normal saline without other compounds served as the control. Tissues from the brain, heart, liver, lung, kidney, and spleen of the mice were harvested at the end of the fifth day. The UVA Research Histology Core Facility conducted paraffin-embedded tissue sectioning and H&E staining.

### BLI of mice with hippocampal luciferases

The process of generating AAVs for neuronal expression of BREP has been detailed previously.^23^ To create analogous viral vectors for expressing Akaluc or Antares, the BREP gene in the pAAV-hSyn-BREP transfer plasmid was substituted with the Akaluc or Antares gene, resulting in the pAAV-hSyn-Akaluc or pAAV-hSyn-Antares transfer plasmid for viral packaging. The titers of the purified viral stocks were quantified using qPCR and were subsequently diluted with Dulbecco’s Phosphate-Buffered Saline (DPBS) to achieve concentrations of 10^14^ GC/mL before use. 1 μL of the diluted virus was bilaterally injected into the hippocampus (relative to Bregma: AP −1.7, ML ± 1.2 and DV −1.5) of 8-week-old BALB/cJ mice via intracranial stereotactic injection at a flow rate of 100 nL/min. Mice were imaged two to three weeks later. A procedure described previously was employed to dissolve DTZ or ETZ in a buffer comprising 25% (w/v) HPCD and 20% (v/v) PEG-400, and 5% (w/v) NaHCO_3_ was additionally added to facilitate the dissolution of ETZ.^23^ Alternatively, the luciferins were simply dissolved in normal saline (9% NaCl). Right before imaging, mice received a 100 μL intravenous injection of the dissolved luciferins. They were subsequently imaged using a UVP BioSpectrum dark box, a Computar Motorized ZOOM lens (M6Z1212MP3), and a Photometrics Evolve 16 EMCCD camera. The lens adjustments were made through the UVP VisionWorksLS software (Version 8.6), with the aperture set to 100% open, zoom at 0%, and focus to 0%. Micro-Manager (Version 2.0.0) was utilized to control the camera for imaging acquisition. The camera analog gain was set to high, and the EM gain was 500. Other parameters included a camera binning of 2 × 2, camera temperature maintained at −70 °C, and exposure time of 100 ms with acquisitions every 30 s. Mice were anesthetized and positioned 23 cm away from the lens without an emission filter. The images were processed using the Fiji version of ImageJ 2.14. Initial background subtraction from the image stacks was conducted with a rolling ball radius set to 100 pixels. Subsequently, a region of interest (ROI) was defined based on the bioluminescence signal from the mouse brain, and the integrated intensity value over the ROI was extracted for further analysis. Despite software-based background subtraction, residual background was noted in the images. To address this, the ROI was relocated away from the mouse brain region to evaluate the residual background, which was then subtracted from the signals to calculate the integrated bioluminescence intensity (area under the curve). The data was plotted and subjected to statistical analysis using GraphPad Prism (Version 8.4.3).

### BLI of mice with liver BREP luciferase

To evaluate sDTZ for deep-tissue imaging in the liver, we developed a pAAV-TBG-BREP transfer plasmid featuring a liver-specific promoter. The AAVs were produced and purified following established protocols.^23^ Subsequently, we administered 100 μL (approximately 1×10^11^ GC/mL) of AAV-TBG-BREP to mice via tail vein injection. After a 3-week period, the mice were subjected to brightness assessment. Luciferins were dissolved in saline and administered intravenously. All parameters for BLI were consistent with those outlined in the previous section.

### BLI of freely moving mice with hippocampal BREP luciferase

BALB/cJ mice expressing hippocampal BREP were prepared following the procedure described above. Before imaging, 100 μL of 25 mM sDTZ was intravenously injected into awake mice. The mice were then positioned in the dark box for BLI. The imaging setup was similar to the one described above, with the addition of a TTL device to regulate the camera and an 850 nm LED light source.

During each imaging cycle, the LED and camera were activated for 15 ms to capture a brightfield image, immediately followed by a 15 ms frame of bioluminescence with the camera on and LED off. The camera binning was adjusted to 8 x 8, and other parameters remained consistent with the settings detailed above. As large data sets were derived and the system was unable to sustain continuous generation of TTL pulses beyond 8.4 seconds, we recorded separate 8.4-second videos at time intervals of 0, 3, 6, 12, and 18 minutes.

### Engineering and *in vitro* characterization of eBRIC

Random mutations were introduced into BRIC via error-prone PCRs,^36^ and the resultant libraries were screened as previously described^23^ except for the following modification. After preparing *E. coli* cell lysates, 5 μL of the supernatant was mixed with 185 μL of a buffer providing 65 nM free Ca^2+^ (30 mM MOPS, 100 mM KCl, 8.75 mM EGTA, 1.25 mM CaEGTA, pH 7.2) or another buffer providing 1.35 μM free Ca^2+^ (30 mM MOPS, 100 mM KCl, 5 mM EGTA, 5 mM CaEGTA, pH 7.2). 10 μL of DTZ solution (500 μM) was dispensed into each well using a reagent injector in a BMG Labtech CLARIOstar Plus microplate reader, resulting in a final DTZ concentration of 25 μM. The bioluminescence spectrum of each well was recorded from 450 to 700 nm with 10 nm intervals. Mutants exhibiting both high brightness and Ca^2+^ responsiveness were chosen for further analysis. Through three rounds of directed evolution, eBRIC was developed. The expression, purification, concentration determination, and *in vitro* characterization of the eBRIC protein were conducted following the established procedures used for BRIC.^23^

### Characterization of eBRIC in HeLa cells

The gene for eBRIC was cloned into pcDNA3 to create pcDNA3-eBRIC, with pcDNA3-BRIC^23^ utilized for comparison. Next, 3 μg of plasmid DNA was used to transfect HeLa cells (obtained from ATCC, Cat. # CCL-2) in each 35-mm culture dish. After overnight incubation at 37 °C in a 5% CO2 incubator, cells were rinsed twice with DPBS (no Ca^2+^ and Mg^2+^), followed by a 20-minute incubation in DPBS before imaging using an inverted Leica DMi8 microscope equipped with a Photometrics Prime 95B Scientific CMOS camera and controlled by Leica LAS X (Version 3.5.7) software. Bioluminescence was initiated by exchanging DPBS with fresh DPBS containing 100 μM DTZ. Imaging parameters included a 40× oil immersion objective lens (NA 1.2), no filter cube, 2×2 camera binning, 1 s exposure with 5 s intervals, camera sensor temperature set at −20 °C, and camera in 12-bit high sensitivity mode. Histamine, dissolved in DPBS, was introduced at a final concentration of 20 μM during the time-lapse imaging. Image processing and data analysis were conducted following established protocols.^23^

### Characterization of eBRIC in primary mouse neurons

The production of AAVs, neuron preparation, transduction, and culture procedures were conducted following established protocols.^23^ Neurons expressing eBRIC and BRIC were assessed on the fifth day post-transduction with the corresponding AAVs. Prior to imaging, the growth medium was substituted with 1.6 mL of luminescence imaging buffer (0.49 mM MgCl_2_, 2 mM CaCl_2_, 0.4 mM MgSO_4_, 0.44 mM KH_2_PO_4_, 5.3 mM KCl, 4.2 mM NaHCO_3_, 0.34 mM Na_2_HPO_4_, 138 mM NaCl, 10 mM HEPES pH 7.2, 15 mM D-glucose, and 0.1 mM sodium pyruvate) supplemented with 100 μM DTZ. During time-lapse imaging, 0.42 mL of high K^+^ stimulation buffer (0.49 mM MgCl_2_, 2 mM CaCl_2_, 0.4 mM MgSO_4_, 0.44 mM KH_2_PO_4_, 143.2 mM KCl, 4.2 mM NaHCO_3_, 0.34 mM Na_2_HPO_4_, 10 mM HEPES pH 7.2, 15 mM D-glucose, and 0.1 mM sodium pyruvate) was applied to depolarize neurons. The imaging setup and data analysis were consistent with the previous section, except for an exposure time of 2 s with 3 s intervals.

### BLI of brain Ca^2+^ dynamics in awake mice

1 μL of AAV-hSyn-eBRIC (5×10^13^ GC/mL) or an equivalent amount of AAV-hSyn-BREP was bilaterally injected into the basolateral amygdala (BLA; coordinates relative to Bregma: AP −3.4, ML ±1.25, DV −4.9)^23^ of 8-week-old BALB/cJ mice. BLI was conducted three weeks post-viral administration. Prior to imaging, each awake mouse received a tail vein injection of 100 μL of sDTZ (25 mM) in normal saline. The mouse was then secured in a Narishige plastic mouse head holder (SRP-AM2) for imaging. BLI was performed using a UVP BioSpectrum dark box, a Computar Motorized ZOOM lens (M6Z1212MP3), and a Photometrics Evolve 16 EMCCD camera. The camera settings included an EM gain of 1000, 8 × 8 binning, an exposure time of 100 ms without intervals, and a sensor temperature of −70 °C. The mice were positioned 27 cm from the lens. Each experimental session consisted of a 100 s acclimation period for the animals, followed by 13 footshock trials. Each trial involved a 0.8-mA electric footshock lasting 1 s, with 40 s intervals between shocks, administered using an A-M Systems 2100 isolated pulse stimulator. Image processing and data analysis were performed according to an established procedure.^23^

## Author Contributions

H.A. and X.T. designed the overall project. X.T. designed and synthesized compounds, performed stability assays, engineered and characterized eBRIC *in vitro* and in cultured cells, constructed plasmids, and prepared viruses. Y.Z performed intracranial viral injection and prepared primary neurons. Y.Z. and X.T. performed animal imaging. Y.Z. performed animal toxicity experiments. X.T. and Y. Z. analyzed data and prepared figures. H.A., X.T. and Y. Z. wrote the manuscript. H.A. provided supervision and obtained funding.

## Acknowledgments

We would like to acknowledge the UVA Biomolecular Magnetic Resonance Facility and Research Histology Core Facility for providing resources and services. Special thanks to Dr. Jing Li for assisting in the toxicology study. The research presented in this publication was supported by the University of Virginia Start-up Fund and National Institutes of Health grant R01EB035430 awarded to HA. ChatGPT was used to paraphrase sentences and rectify grammatical errors.

